# Intrinsic and tumor-induced JAK/STAT signaling regulate developmental timing by the *Drosophila* prothoracic gland

**DOI:** 10.1101/2021.06.09.447673

**Authors:** Xueya Cao, José Carlos Pastor-Pareja

**Affiliations:** School of Life Sciences, Tsinghua University, Beijing, China; Tsinghua-Peking Center for Life Sciences, Beijing, China

**Keywords:** Metamorphosis, ring gland, ecdysone, tissue damage, inflammation, SUMOylation

## Abstract

Development involves tightly paced, reproducible sequences of events, yet it must adjust to conditions external to it, such as resource availability and organismal damage. A major mediator of damage-induced immune responses in vertebrates and insects is JAK/STAT signaling. At the same time, JAK/STAT activation by the *Drosophila* Upd cytokines is pleiotropically involved in normal development of multiple organs. Whether inflammatory and developmental roles of JAK/STAT intersect is unknown. Here, we show that JAK/STAT is active during development of the prothoracic gland (PG), the organ that controls metamorphosis onset through ecdysone production. Reducing JAK/STAT signaling decreased PG size and slightly advanced metamorphosis. Conversely, JAK/STAT hyperactivation, achieved through overexpression of pathway components or SUMOylation loss, caused PG hypertrophy and metamorphosis delay. Interestingly, tissue damage and tumors, known to secrete Upd cytokines, also activated JAK/STAT in the PG and delayed metamorphosis. Finally, we show that expression of transcription factor Apontic, a JAK/STAT target in the PG, recapitulates PG hypertrophy and metamorphosis delay. JAK/STAT damage signaling, therefore, regulates metamorphosis onset at least in part by coopting its developmental role in the PG.

**Summary statement:** Damage signaling from tumors mediated by JAK/STAT-activating Upd cytokines delays the *Drosophila* larva-pupa transition through cooption of a JAK/STAT developmental role in the prothoracic gland.

## INTRODUCTION

The development of organisms involves internally paced, reproducible sequences of molecular and cellular events. In the development of holometabolous insects, a larval stage with little resemblance to the adult transitions to a non-feeding pupal stage during which metamorphosis takes place. This life cycle strategy arose 350 million years ago in the Carboniferous period and led to the amazing radiation of the four most successful orders of insects: coleopterans, lepidopterans, hymenopterans and dipterans (Truman and Riddiford, 2019). The temporal regulation of the larva-pupa transition depends on hormones produced by the neuroendocrine system, chief among them the steroid ecdysone, secreted by the cells of the prothoracic gland (Tennessen and Thummel, 2011; Yamanaka et al., 2013). The prothoracic gland (PG) is part of the ring gland, a tripartite organ situated anterior to the brain and additionally consisting of the corpus allatum and the corpora cardiaca (Fig. 1A). The cells of the PG and corpus allatum have an ectodermal origin in tracheal primordia of the embryonic head, whereas the corpora cardiaca derive from mesoderm (Sanchez-Higueras et al., 2014). Studies in the fruit fly *Drosophila melanogaster*, a dipteran, have shown that multiple molecular signals influence ecdysone production by the PG, including PTTH, insulin and TOR, Hippo, TGFβ, EGF, DILP8, nitric oxide, the circadian clock, ecdysone itself, tyramine, serotonin and Hh (Texada et al., 2020). Their coordinated actions on PG cells integrate developmental inputs and environmental cues to determine the timing of ecdysone synthesis and release. Although usually less pronounced, similar hormonally controlled developmental transitions are common in animal development. Indeed, puberty transition and insect metamorphosis might share a common Urbilaterian ancestry (Barredo et al., 2021). In general, hormonal control of developmental timing and transitions maximizes organismal fitness by ensuring development that is reproducible, robust and adjusted to conditions external to it, such as resource availability, environmental insults and damage to the organism.

**Figure 1.**
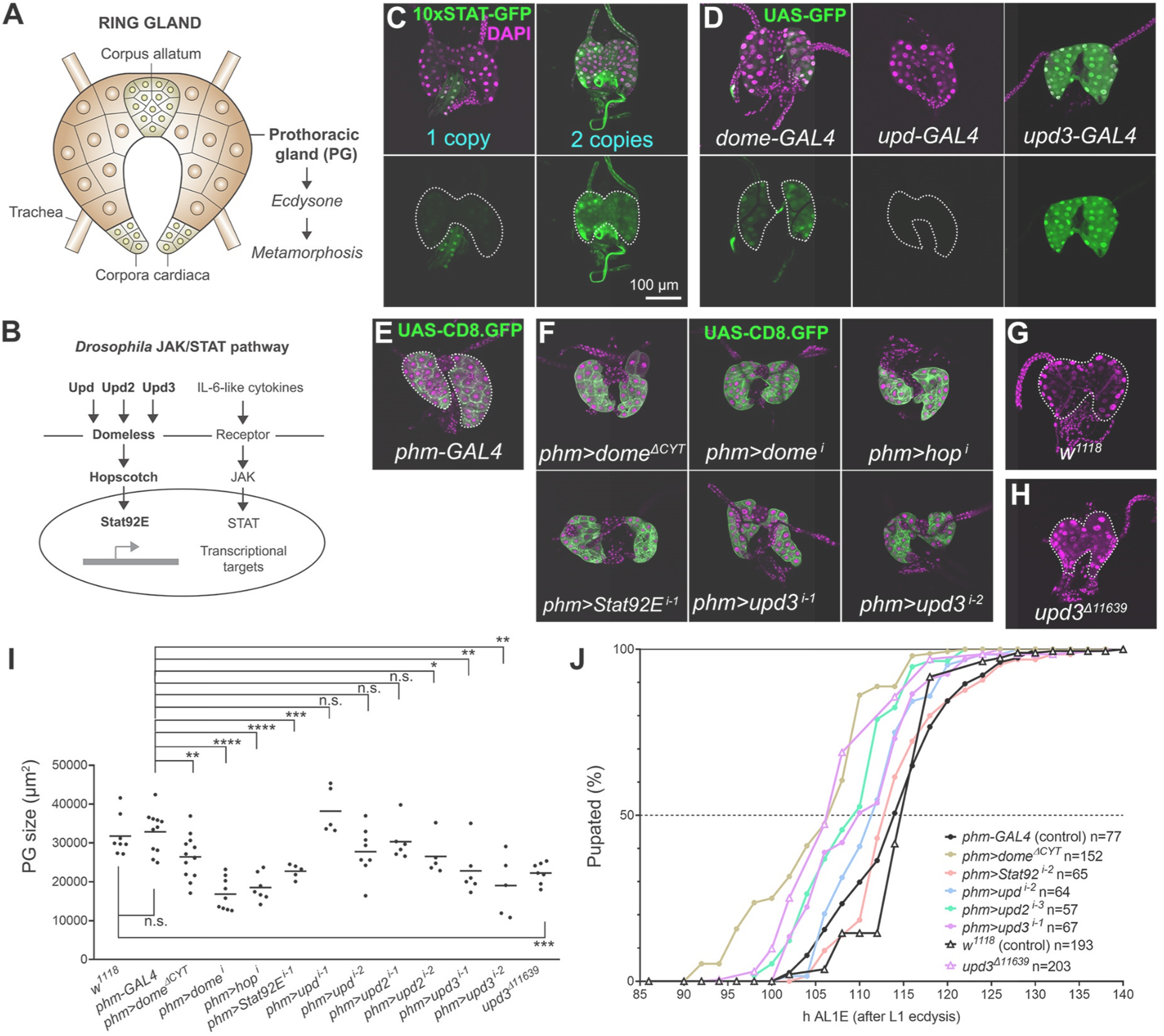
Intrinsic JAK/STAT signaling modulates PG growth and metamorphosis. (A) Schematic drawing of the ring gland of the third instar *Drosophila* larva. The prothoracic gland (PG), producing ecdysone, is the largest part of this composite neuroendocrine organ. (B) Components of the JAK/STAT signaling pathway in *Drosophila*. (C) Confocal images of ring glands dissected from wandering third instar (wL3) larvae containing one (left) or two (right) copies of the JAK/STAT activity reporter 10xSTAT-GFP (Ekas et al., 2006) (green, separate channel in lower subpanels). Dotted lines represent PG outline. Nuclei stained with DAPI (magenta). (D) Ring glands from wL3 larvae expressing GFP (green, separate channel in lower subpanels) under control of GAL4 transcriptional reporters for genes encoding receptor Domeless (left), and cytokines Upd (center) and Upd3 (right). Dotted lines represent PG outline. Nuclei stained with DAPI (magenta). (E) Ring gland from a control wL3 larva expressing membrane GFP (CD8.GFP, green) under control of PG-specific *phm-GAL4*. Dotted lines represent PG outline. Nuclei stained with DAPI (magenta). (F) Ring glands from wL3 larvae expressing dominant negative Dome^ΔCYT^ or knocking down *dome, hop, Stat92E* and *upd3* in the PG under control of *phm-GAL4*. Knock down of *upd3* in *phm>upd3*^*i-1*^ and *phm>upd3*^*i-2*^ experiments employs different RNAi transgenes (see Table S1 for detailed genotypes in these and all other experiments throughout the manuscript). *phm-GAL4*-driven CD8.GFP in green. Nuclei stained with DAPI (magenta). (G) Ring gland from a *w*^*1118*^ control wL3 larva. Dotted lines represent PG outline. Nuclei stained with DAPI (magenta). (H) Ring gland from a null *upd3*^*Δ11639*^ mutant wL3 larva. Dotted lines represent PG outline. Nuclei stained with DAPI (magenta). (I) Quantification of wL3 PG size in *phm-GAL4* and *w*^*1118*^ control larvae, larvae expressing dominant negative Dome^ΔCYT^ or knocking down *dome, hop, Stat92E, upd, upd2* and *upd3* under control of *phm-GAL4*, and null *upd3*^*Δ11639*^ mutant larvae. Each dot plots PG size measured as the area occupied by the PG in images of ring glands like those in panels E-H. Horizontal lines represent mean values. Conducted tests were Mann-Whitney tests, except for comparison of both controls and for *phm>dome*^*ΔCYT*^, *phm> dome*^*i*^ and *phm>upd*^*i-1*^ (two-tailed t-tests). Significance of differences in statistical tests is reported in all figures as follows: p<0.0001 (****), p<0.001 (***), p<0.01 (**), p<0.05 (*), and p>0.05 (n.s., not significant). (J) Pupation time in *phm-GAL4* and *w*^*1118*^ controls, and in the indicated JAK/STAT loss of function conditions. Dot-connecting lines plot the accumulated percentage of pupated larvae through time. Time of pupation was computed as hours after L1 ecdysis (see Materials and methods). Number of animals examined is reported in the graph.

Tissue damage signaling activated by mechanical wounding (Bryant, 1971; Díaz-García and Baonza, 2013) or high levels of cell death (Akai et al., 2021; Hackney et al., 2012; Simpson and Schneiderman, 1975; Smith-Bolton et al., 2009; Stieper et al., 2008) is well known to delay the onset of metamorphosis. Damage-induced extension of the larval period may have evolved to allow for more complete healing and regeneration before transition to the next developmental stage occurs (Hariharan and Serras, 2017). Tumorous growth perturbations, sharing tissue damage inflammatory mechanisms with wounds (Pastor-Pareja et al., 2008; Pastor-Pareja and Xu, 2013), delay or completely inhibit the larva-pupa transition as well (Menut et al., 2007; Pagliarini and Xu, 2003; Sehnal and Bryant, 1993). One of the signals mediating damage-induced metamorphosis delay is the biosynthesis of retinoids (Halme et al. 2010). Through poorly understood mechanisms, retinoids inhibit production in the central nervous system of prothoracicotropic hormone (PTTH), required for ecdysone production by the PG. Another signal acting on PG ecdysone production through inhibition of PTTH synthesis is Dilp8, a member of the insulin-like/relaxin family of peptide hormones. Dilp8 is produced by damaged tissues and tumors downstream of the c-Jun N-terminal kinase (JNK) pathway (Colombani et al., 2012; Garelli et al., 2012). Autocrine JAK/STAT signaling, activated in wounds and tumors by JNK-induced expression of Upd cytokines (Pastor-Pareja et al., 2008; Wu et al., 2010), has been reported to enhance local Dilp8 production by the damaged tissue (Katsuyama et al., 2015). In the tissue damage response, however, JAK/STAT-activating Upd cytokines have been shown to act not just locally, driving damage-induced regenerative growth (Katsuyama et al., 2015; La Fortezza et al., 2016; Santabarbara-Ruiz et al., 2015; Wu et al., 2010), but also systemically. Indeed, damaged-induced Upd cytokines mediate an innate immune response that induces proliferation of circulating macrophages and amplifies itself through additional Upd cytokine expression in the fat body (Pastor-Pareja et al., 2008). Furthermore, Upd cytokines have been shown to act systemically as well for inter-organ communication of nutritional status in the adult (Rajan and Perrimon, 2012). Additional, non-local roles of damage-induced Upd cytokines in developmental timing are therefore possible.

The JAK/STAT (Janus kinase and signal transducer and activator of transcription) signaling pathway (Fig. 1B), is a major mediator of innate immune responses in vertebrates and insects alike (Agaisse and Perrimon, 2004; Lemaitre and Hoffmann, 2007). In *Drosophila*, JAK/STAT signaling is activated by three highly similar Interleukin-6-related cytokine ligands: Unpaired (Upd) (Harrison et al., 1998), Upd2 (Gilbert et al., 2005; Hombría et al., 2005) and Upd3 (Agaisse et al., 2003). Upd cytokines bind to the transmembrane receptor Domeless (Dome) (Brown et al., 2001; Chen et al., 2002), inducing activation of the receptor-associated Janus kinase (JAK) homolog Hopscotch (Hop) (Binari and Perrimon, 1994). Activated Hop in turn phosphorylates STAT transcription factor homolog Stat92E (Hou et al., 1996; Yan et al., 1996), which then translocates as a dimer to the nucleus to activate expression of target genes (Arbouzova and Zeidler, 2006; Rawlings et al., 2004; Zeidler and Bausek, 2013). In addition to its immune roles, JAK/STAT activation by Upd cytokines is pleiotropically involved in normal development and differentiation of multiple non-immune organs and cell types in *Drosophila*. Examples of non-immune contexts where JAK/STAT signaling is known to play a developmental role include segmental patterning (Binari and Perrimon, 1994), tracheae (Brown et al., 2001), border cells of the ovarian follicle (Beccari et al., 2002; Silver and Montell, 2001), the eye (Zeidler et al., 1999), the wing (Recasens-Alvarez et al., 2017), the optic lobe (Yasugi et al., 2008), and multiple stem cell niches (Herrera and Bach, 2019). A systemic JAK/STAT immune response, such as the tissue damage response, may therefore influence development of multiple tissues. Whether the inflammatory and developmental functions of JAK/STAT signaling intersect, however, has not been investigated.

Here, we studied the role of JAK/STAT signaling in the PG. We found that JAK/STAT signaling is active during normal PG development. Reduced JAK/STAT signaling decreased average PG size and slightly advanced metamorphosis, while pathway hyperactivation caused PG hypertrophy and delayed the larva/pupa transition. Tissue damage and tumors, known to secrete Upd cytokines, also led to JAK/STAT activation in the PG and metamorphosis delay. JAK/STAT effects on the PG, our results indicate, are mediated at least in part by downstream expression of Apontic. Altogether, our experiments reveal that damage signaling by Upd cytokines regulates the onset metamorphosis by directly activating JAK/STAT signaling in the PG, thus coopting a developmental role of JAK/STAT therein.

## RESULTS

### Intrinsic JAK/STAT signaling modulates PG growth and metamorphosis

To investigate a possible role of JAK/STAT signaling in ring gland development and developmental timing, we examined expression of 10xSTAT-GFP, a reporter of JAK/STAT signaling in which GFP expression is driven by 10 copies of the STAT binding sequence in the promoter of JAK/STAT target *Socs36E* (Ekas et al., 2006). In ring glands dissected from wandering third instar larvae (wL3), we were able to detect expression of the STAT-GFP reporter in the prothoracic gland (PG), increased when two copies of the reporter were present in homozygous flies (Fig. 1C). In addition, we found that reporters *dome-GAL4* and *upd3-GAL4*, transcriptional reporters for the expression of *domeless*, encoding the JAK/STAT receptor, and *unpaired3*, encoding one of the three JAK/STAT-activating, Interleukin-6-related cytokines present in *Drosophila*, were expressed in the PG as well (Fig. 1D), further suggesting that JAK/STAT signaling is active in the PG. To test a role of JAK/STAT signaling in the PG, we examined ring glands in different conditions affecting JAK/STAT signaling. Knock down of the genes encoding receptor Domeless, the kinase Hopscotch and transcription factor Stat92E under control of PG-specific *phm-GAL4*, as well as expression of dominant negative Domeless (Dome^ΔCYT^), resulted in ring glands that were in average smaller than controls (Fig. 1E and F, quantified in Fig. 1J). Among the three JAK/STAT ligands, knock down of *upd3* with two different RNAi transgenes showed a consistent decrease on ring gland size (Fig. 1E, F and J), while knock down of *upd* and *upd2* had not significant or less significant effects. Furthermore, null *upd3* mutants (Osman et al., 2012) also showed significantly reduced PG size (Fig. 1G-I). Because the PG has a major role in regulating developmental timing, we next investigated the effect of JAK/STAT signaling reduction in the timing of the larva/pupa transition. To do that, we recorded the time of pupation of animals after L1 ecdysis (embryo/L1 transition). Compared to controls, we observed that the time of the larva/pupa transition was advanced in different JAK/STAT loss of function conditions, with *upd3* null mutants and Dome^ΔCYT^ expression in the PG showing the clearest such advanced pupation effects (Fig. 1K). In all, these results indicate that intrinsically-activated JAK/STAT signaling is active in the PG, where it contributes to its development and modulates the timing of the larva/pupa transition.

### JAK/STAT hyperactivation in the PG delays metamorphosis

To further investigate the role of JAK/STAT signaling in the PG, we aimed at increasing JAK/STAT activity and studying its effects on PG development and metamorphosis onset. To do that, we overexpressed JAK/STAT cytokines Upd, Upd2 and Upd3 in the PG under control of *phm-GAL4*. We found that expression of all three cytokines showed similar ability to activate JAK/STAT signaling, as evidenced by highly increased expression of the 10xSTAT-GFP activity reporter (Fig. 2A, quantified in Fig. 2B). In addition to this, we observed that larvae expressing JAK/STAT cytokines in their PG did not pupate and became giant larvae (Fig. 2C). The PG in these larvae, and in larvae overexpressing Dome and Hop in their PG, became highly enlarged and abnormal, with some cells presenting a large degree of vacuolation (Fig. 2D and E), suggestive of a degenerative or autolytic process. PG hypertrophy was accompanied by a spectacular increase in nuclear size and ploidy of these cells estimated by their DNA content, while average PG ploidy was reduced upon knock down of *Stat92E* (Fig. 2F). The number of PG nuclei in JAK/STAT loss (*phm>Stat92E*^*i*^) and gain (*phm>upd*^*OE*^) conditions, however, did not change with respect to the wild type (Fig. 2G). Consistent with their inability to undergo larva/pupa transition, quantitative real time PCR showed that the PG of animals overexpressing Upd expressed reduced levels of *disembodied* (*dib*), *neverland* (*nvd*), *phantom* (*phm*) and *shadow* (*sad*), encoding Halloween group enzymes, involved in the synthesis of ecdysone (Fig. 2H, results of three biological replicates). All these results support a positive role of JAK/STAT signaling in PG cell growth and endoreplication. At the same time, they indicate that while basal levels of JAK/STAT signaling counter premature metamorphosis, JAK/STAT hyperactivation has the opposite effect and prevents pupation.

**Figure 2.**
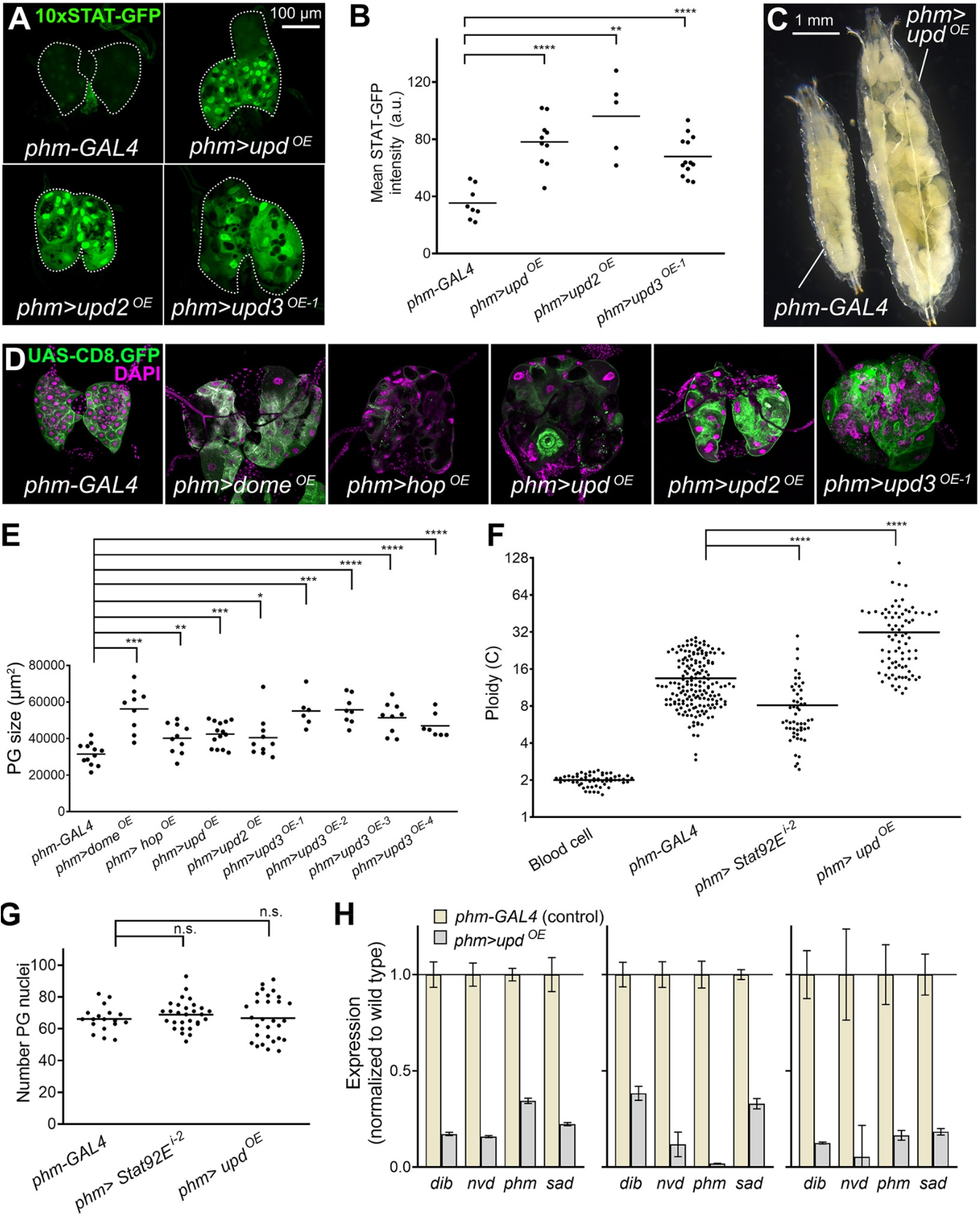
JAK/STAT hyperactivation causes PG hypertrophy and delayed metamorphosis. (A) Expression of JAK/STAT activity reporter 10xSTAT-GFP (green, separate channel in lower subpanels) in the PG of control *phm-GAL4* wL3 larvae and giant larvae overexpressing *upd, upd2*, and *upd3* under control of *phm-GAL4*. Dotted lines represent PG outline. (B) Quantification of 10xSTAT-GFP intensity in PG of the larvae overexpressing *upd, upd2*, and *upd3* under *phm-GAL4* control. Each dot plots mean GFP intensity (total fluorescence intensity divided by area) in the PG measured in images like those in panel A. Horizontal lines represent mean values. Significance of differences in two-tailed t-tests (*phm>upd*^*OE*^ and *phm>upd3*^*OE*^) and Mann-Whitney tests (*phm>upd2*^*OE*^) is reported. (C) Control *phm-GAL4* wL3 larva (left) and larva overexpressing *upd* under *phm-GAL4* control in the PG (*phm>upd*^*OE*^, right). Unable to pupate, *phm>upd*^*OE*^ L3 larvae indefinitely extend their larval period and become giant larvae. (D) Ring glands from a control *phm-GAL4* wL3 larva and from L3 giant larvae overexpressing *dome, hop, upd, upd2* and *upd3* in the PG under control of *phm-GAL4. phm-GAL4*-driven CD8.GFP in green. Nuclei stained with DAPI (magenta). (E) Quantification of PG size in wL3 *phm-GAL4* control larvae and in giant larvae overexpressing JAK/STAT pathway components in the PG under control of *phm-GAL4* as indicated. Each dot plots PG size measured from images like those in panel D. Horizontal lines represent mean values. Significance of differences in statistical tests is reported. Conducted tests were two-tailed t-tests (*phm>hop*^*OE*^, *phm>upd*^*OE*^, *phm>upd3*^*OE-2*^ and *phm>upd3*^*OE-3*^), two-tailed t-tests with Welch’s correction (*phm>dome*^*OE*^) and Mann-Whitney tests (*phm>upd2*^*OE*^, *phm>upd3*^*OE-1*^ and *phm>upd3*^*OE-4*^). (F) Quantification of ploidy in PG nuclei of control *phm-GAL4* wL3 larvae, *phm>Stat92E*^*i-2*^ wL3 larvae and giant *phm>upd*^*OE-1*^ larvae. Each dot plots ploidy of a nucleus calculated based on DNA content (DAPI intensity) integrated from confocal stacks of the whole PG and with reference to DNA content in diploid blood cells (see Materials and Methods). n=58, 172, 53 and 80 nuclei, respectively. Horizontal lines represent mean values. Significance of differences in Mann-Whitney tests is reported. (G) Number of nuclei in *phm-GAL4* wL3 larvae, *phm>Stat92E*^*i-2*^ wL3 larvae and giant *phm>upd*^*OE*^ larvae. Each point represents the number of PG nuclei in one ring gland as counted manually from confocal stacks of the whole PG. n=18, 29 and 29, respectively. Differences were not significant in two-tailed t-tests (*phm>Stat92E*^*i-2*^) and Mann-Whitney tests (*phm>upd*^*OE*^). (H) Quantification of expression of Halloween genes of the ecdysone synthesis pathway by qRT-PCR. Expression levels relative to *rpL23* were normalized to the wild type *phm-GAL4* control. Three biological replicates are shown, each of them with three technical replicates (error bars represent SD).

We were intrigued by the JAK/STAT hyperactivation PG phenotype combining enlargement and partial tissue destruction. We found in the literature a strikingly similar phenotype caused by loss of function of *smt3*, encoding the *Drosophila* homolog of Ubiquitin-like posttranslational modifier SUMO (Talamillo et al., 2008). Furthermore, the E3 SUMO ligase Su(var)2-10, also known as PIAS (Protein Inhibitor of Activated STATs), regulates STAT activity in *Drosophila* and humans (Betz et al., 2001; Grönholm et al., 2010). We found that knock down of *smt3, Su(var)2-10* and other components of the *Drosophila* SUMOylation cascade strongly increased JAK/STAT activity in the ring gland, evidenced by increased 10xSTAT-GFP reporter expression (Fig. 3A, quantified in Fig. 3B). Extreme PG hypertrophy was observed with RNAi transgenes targeting *Su(var)2-10* (Fig. 3A, quantified in Fig. 3C), producing in addition inhibition of the larva/pupa transition (Fig. 3D). Confirming that SUMOylation affects PG development through JAK/STAT hyperactivation, knock down of *Stat92E* strongly suppressed JAK/STAT activity and hypertrophy in the PG induced by *Su(var)2-10* knock down (Fig. 3E, quantified in Fig. 3F and 3G). Furthermore, *Stat92E* knock down allowed development of these animals into adults (Fig. 3H and I). These results indicate that JAK/STAT activity in the PG is negatively regulated by the SUMOylation cascade and confirm the potential of the JAK/STAT pathway to influence PG development and timing of metamorphosis.

**Figure 3.**
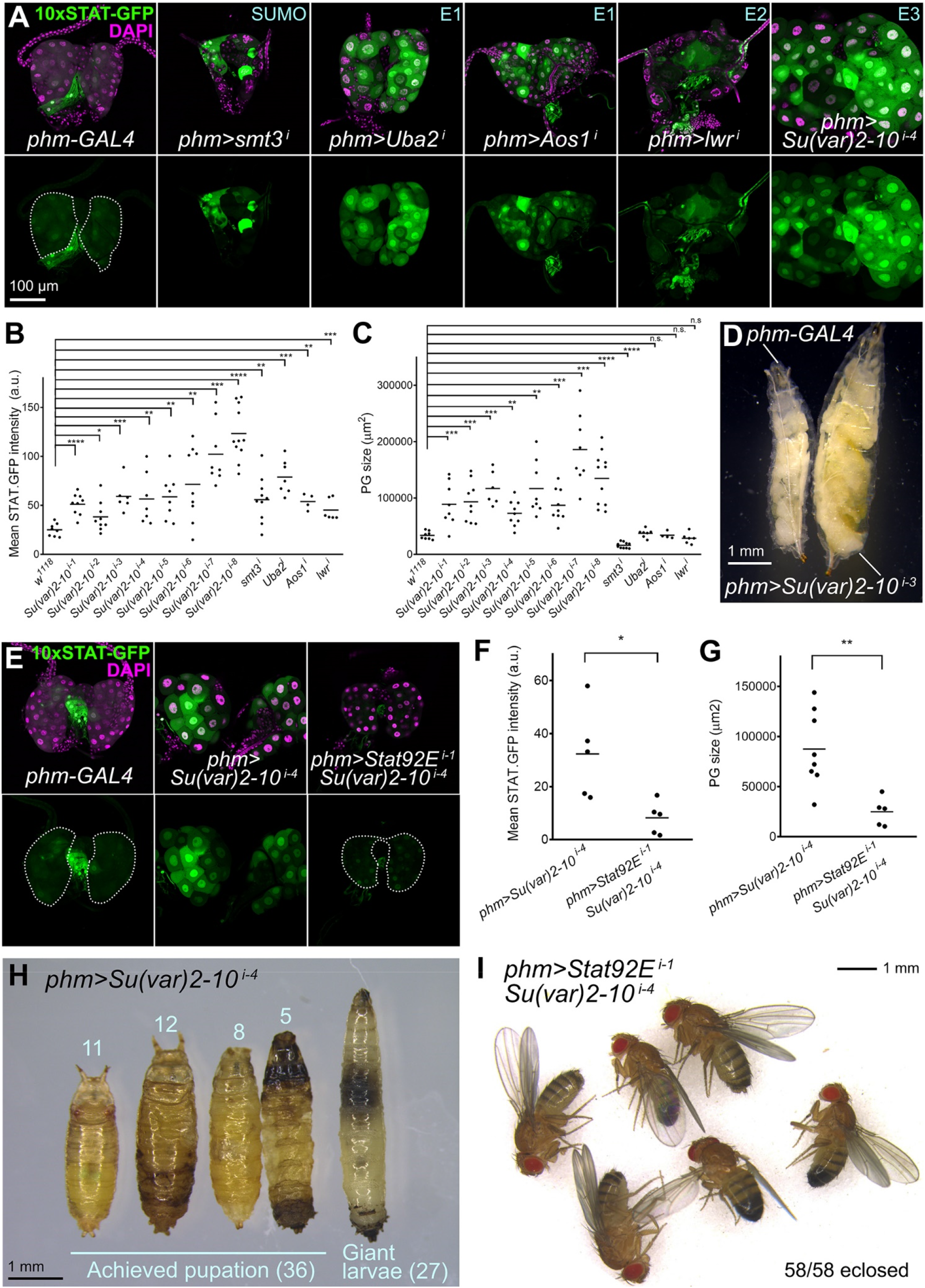
SUMOylation prevents JAK/STAT hyperactivation during normal PG development. (A) Expression of JAK/STAT activity reporter 10xSTAT-GFP (green, separate channel in lower subpanels) in the PG of control *phm-GAL4* wL3 larvae and giant larvae where expression of SUMOylation pathway components has been knocked down under control of *phm-GAL4*. Dotted lines represent PG outline. Nuclei stained with DAPI (magenta). (B) Quantification of 10xSTAT-GFP intensity in PG of control *phm-GAL4* wL3 larvae and giant larvae where expression of SUMOylation pathway components has been knocked down under control of *phm-GAL4*. Each dot plots mean GFP intensity in the PG measured in images like those in panel A. Horizontal lines represent mean values. Significance of differences in statistical tests is reported. Conducted tests were two-tailed t-tests with Welch’s correction except for *phm>Su(var)2-10*^*i-1*^ (two-tailed t-test), *phm>Su(var)2-10*^*i-3*^, *phm>su(var)2-10*^*i-4*^, *phm>Uba2*^*i*^, *phm>Aos1*^*i*^ and *phm>lwr*^*i*^, (Mann-Whitney tests). (C) Quantification of PG size in control *phm-GAL4* wL3 larvae and giant larvae where expression of SUMOylation pathway components has been knocked down under control of *phm-GAL4*. Each dot plots PG size measured from images like those in panel A. Horizontal lines represent mean values. Significance of differences in statistical tests is reported. Conducted tests were two-tailed t-tests with Welch’s correction except for *phm>smt3*^*i*^ (two-tailed t-test), and *phm>Su(var)2-10*^*i-3*^, *phm>Uba2*^*i*^, *phm>Aos1*^*i*^ and *phm>lwr*^*i*^ (Mann-Whitney tests). (D) Control *phm-GAL4* wL3 larva (left) and giant *phm>Su(var)2-10*^*i-3*^ larva (right). (E) Expression of JAK/STAT activity reporter 10xSTAT-GFP (green, separate channel in lower subpanels) in the PG of control *phm-GAL4* wL3 larvae (left) and larvae where expression of *Su(var)2-10* (center) and both *Su(var)2-10* and *Stat92E* (right) has been knocked down under control of *phm-GAL4*. Dotted lines represent PG outline. Nuclei stained with DAPI (magenta). (F) Quantification of 10xSTAT-GFP intensity in PG of wL3 larvae where expression of *Su(var)2-10* and both *Su(var)2-10* and *Stat92E* have been knocked down under control of *phm-GAL4*. Each dot plots PG size measured from images like those in panel E. Horizontal lines represent mean values. Difference was significant in a Mann-Whitney test. (G) Quantification of PG size in wL3 larvae where expression of *Su(var)2-10* and both *Su(var)2-10* and *Stat92E* have been knocked down under control of *phm-GAL4*. Each dot plots mean GFP intensity in the PG measured in images like those in panel E. Horizontal lines represent mean values. Difference was significant in a Mann-Whitney test. (H) Defective pupation in *phm>Su(var)2-10*^*i-4*^ larvae. Number of animals unable to pupate (giant larvae) and exhibiting pupation defects is indicated. Even if pupation was achieved, adults did not eclose. (I) *phm>Su(var)2-10*^*i-4*^ where *Stat92E* is additionally knocked down reach adulthood.

### Tumors and tissue damage activate PG-extrinsic JAK/STAT signaling to delay metamorphosis

Upd cytokines are abundantly produced in innate immune responses (Agaisse and Perrimon, 2004). Of note, Upd cytokines produced by tumors or wounds have been previously shown to activate JAK/STAT signaling in the fat body and blood cells, causing a systemic response through fat body amplification of cytokine production (Pastor-Pareja et al., 2008). Since we had found that JAK/STAT activity regulated pupation time in the PG, we hypothesized that extrinsically produced Upd cytokines could act in the PG similar to intrinsically expressed ones. To test this, we overexpressed cytokines Upd1, Upd2 and Upd3 outside the PG under control of *Cg-GAL4*, expressed in the fat body and circulating blood cells of the hemolymph-filled body cavity (Fig. 4A). In all three cases, we found that PG-extrinsic expression of Upd cytokines was capable of inducing high levels of 10xSTAT-GFP reporter expression in the PG (Fig. 4B, quantified in Fig. 4C). Furthermore, these animals showed absent or delayed metamorphosis (Fig. 4D and E). These results, importantly, show that Upd cytokines produced outside of the PG can activate JAK/STAT signaling and produce developmental delay, similar to that observed when Upd cytokines were overexpressed by PG cells or when activity of the pathway was manipulated inside the PG.

**Figure 4.**
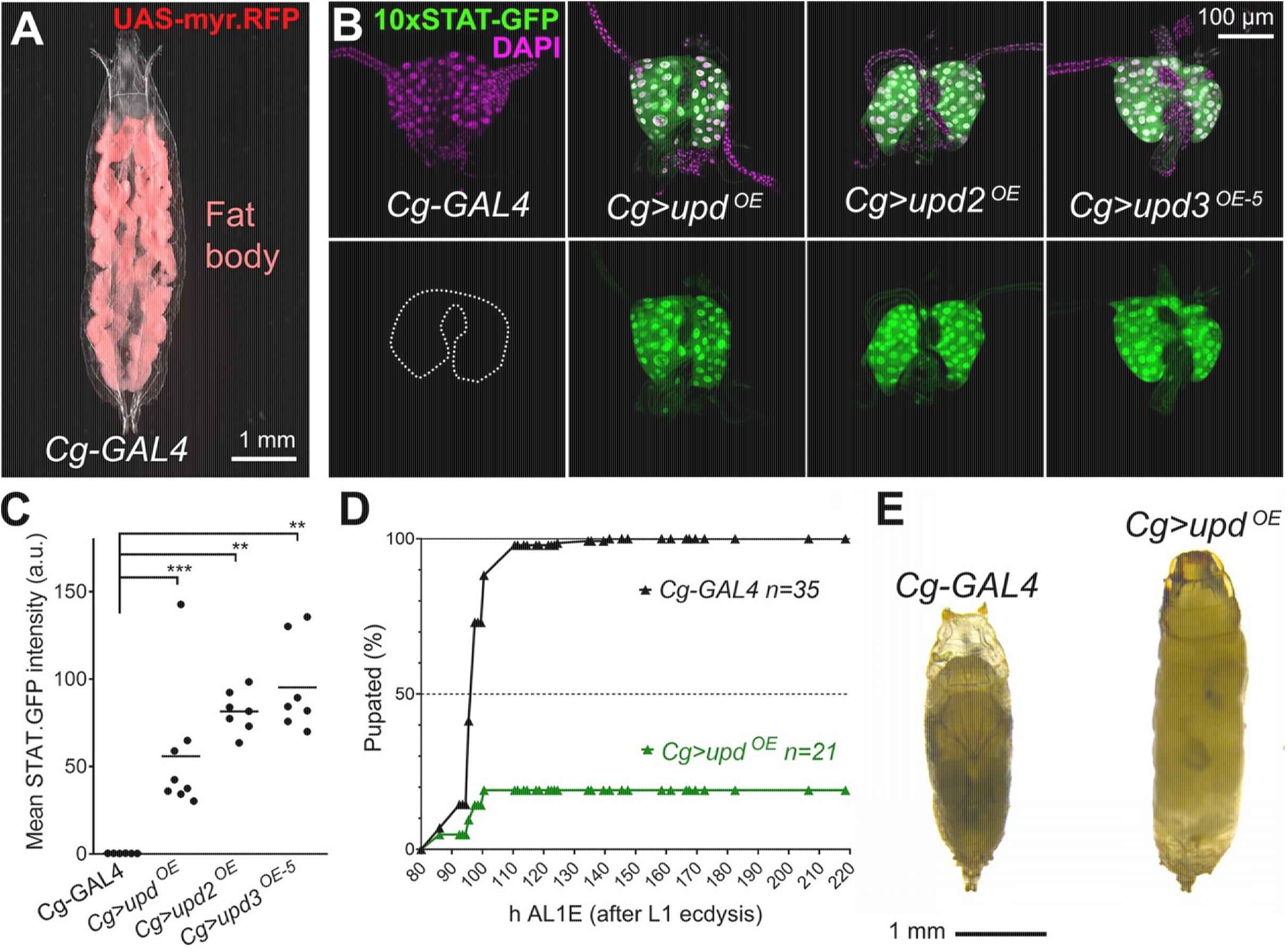
Extrinsically-produced Upd cytokines can activate JAK/STAT signaling in the PG. (A) wL3 larva showing myr.RFP expression in the fat body (red) driven by *Cg-GAL4*. White light and red fluorescence images have been superimposed. (B) Expression of JAK/STAT activity reporter 10xSTAT-GFP (green, separate channel in lower subpanels) in the PG of control *Cg-GAL4* wL3 larvae and giant larvae overexpressing *upd, upd2* and *upd3* in the fat body under control of Cg-GAL4. Dotted lines represent PG outline. Nuclei stained with DAPI (magenta). (C) Quantification of 10xSTAT-GFP intensity in PG of control *Cg-GAL4* wL3 larvae and giant larvae overexpressing *upd, upd2* and *upd3* in the fat body under control of Cg-GAL4. Each dot plots mean GFP intensity in the PG measured in images like those in panel B. Horizontal lines represent mean values. Significance of differences in Mann-Whitney tests is reported. (D) Pupation time in *Cg-GAL4* control and *Cg>upd*^*OE*^ animals. Dot-connecting lines plot the accumulated percentage of pupated larvae through time. (E) Control *Cg-GAL4* pupa (left) and pupa of an animal overexpressing *upd* under control of *Cg-GAL4* in the fat body (right). Even if pupation is achieved, no adults eclose from *Cg>upd*^*OE*^ pupae.

Having shown that extrinsically produced Upd cytokines can activate JAK/STAT signaling in the PG, we asked whether tissue damage conditions elevating systemic Upd cytokine levels induced JAK/STAT activity in the PG. We found that puncture wounding of the larval epidermis as well as neoplastic tumors in *scribbled* (*scrib*) mutants increased JAK/STAT activity in the PG (Fig. 5A, quantified in Fig. 5B), suggesting that upon tissue damage, JAK/STAT activation in the PG could contribute to metamorphosis delay in these conditions. To test this, we employed a mosaic model of clonally-induced tumors in the wing imaginal discs. In this model, large tumors formed by *scribbled warts* (*scrib wts*) homozygous double mutant cells are generated through mitotic recombination in a wild type animal (Fig. 5C). Despite pupation delay by *scrib wts* wing disc tumors, all animals in the end are capable of undergoing metamorphosis. In this tumor model, we found that activation of JAK/STAT signaling in the PG was decreased by expression of dominant negative receptor Dome^ΔCYT^ (Fig. 5D, quantified in Fig. 5E). Furthermore, Dome^ΔCYT^ expression in the PG partially rescued pupation delay in tumor-bearing animals (Fig. 5F, four repeats of the same experiment). Similarly, Dome^ΔCYT^ expression in the PG partially rescued metamorphosis delay in an experiment where damage was induced by heating mid third instar larvae at 39^0^C for 2.5 hours (Fig. 5G). Altogether, these results show that JAK/STAT activation in the PG extrinsically induced by tumors and tissue damage contributes to delaying the larva/pupa developmental transition.

**Figure 5.**
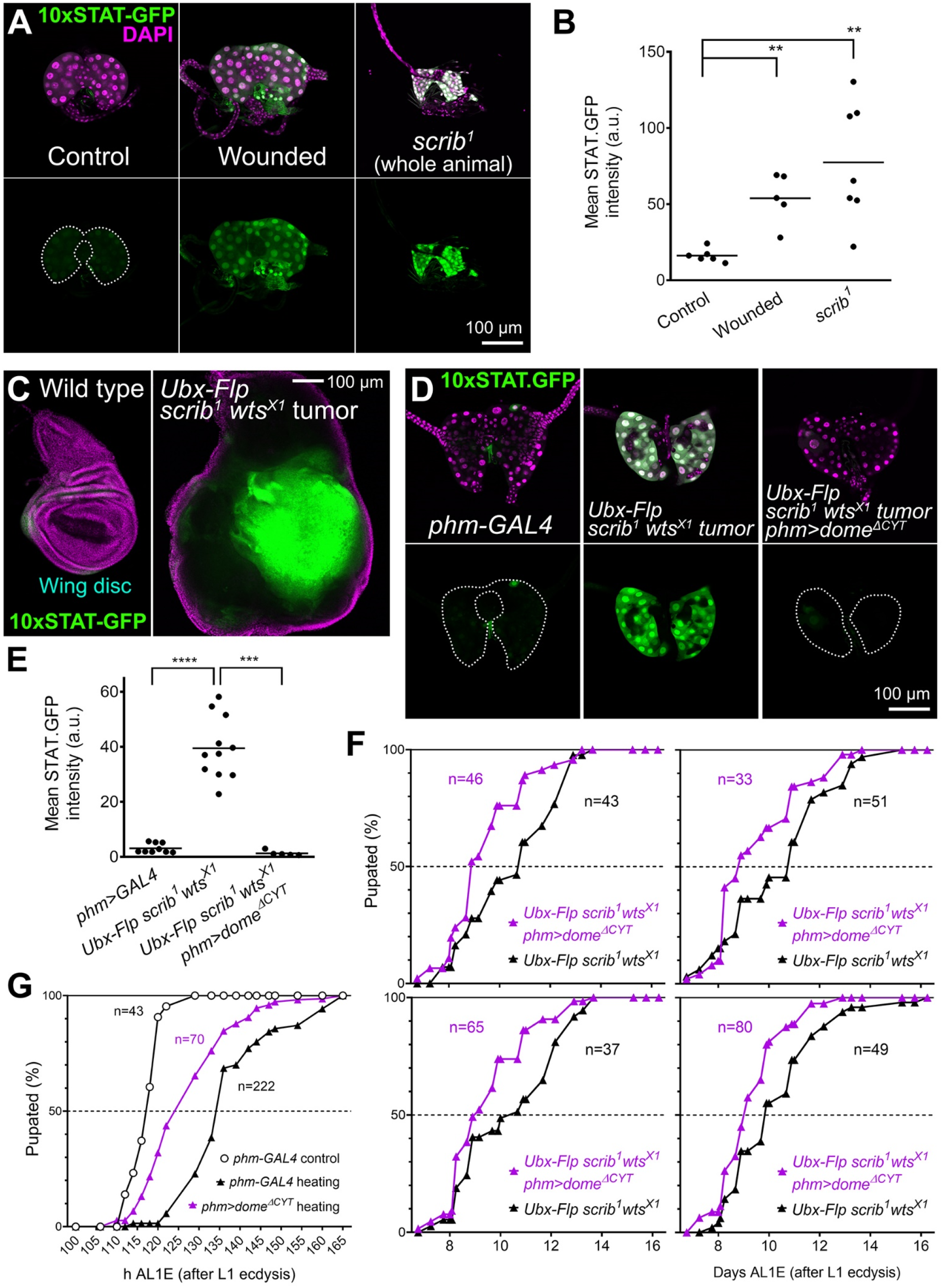
Tumors and tissue damage activate PG-extrinsic JAK/STAT signaling to delay metamorphosis. (A) Expression of JAK/STAT activity reporter 10xSTAT-GFP (two copies, green, separate channel in lower subpanels) in the PG of control (left) and puncture-wounded wL3 larvae (center), and of tumor-containing *scrib*^*1*^ giant larvae. Dotted lines represent PG outline. Nuclei stained with DAPI (magenta). (B) Quantification of 10xSTAT-GFP intensity in PG of control and puncture-wounded wL3 larvae (center), and of tumor-containing *scrib*^*1*^ giant larvae. Each dot plots mean GFP intensity in the PG measured in images like those in panel A. Horizontal lines represent mean values. Differences in Mann-Whitney tests were significant. (C) Expression of JAK/STAT activity reporter 10xSTAT-GFP (green, separate channel in lower subpanels) in a control wL3 wing imaginal disc (left) and in a wing disc from a wL3 larva containing *scrib*^*1*^ *wts*^*X1*^ tumor clones induced through Ubx-Flp-driven mitotic recombination (right). Nuclei stained with DAPI (magenta). (D) Expression of JAK/STAT activity reporter 10xSTAT-GFP (green, separate channel in lower subpanels) in the wL3 PG of control *phm-GAL4* larvae (left), larvae with *scrib*^*1*^ *wts*^*X1*^ tumor clones in the wing disc (center) and larvae with *scrib*^*1*^ *wts*^*X1*^ wing tumors additionally expressing Dome^ΔCYT^ in the PG under *phm-GAL4* control (right). Dotted lines represent PG outline. Nuclei stained with DAPI (magenta). (E) Quantification of 10xSTAT-GFP intensity in the wL3 PG of control *phm-GAL4* larvae, larvae with *scrib*^*1*^ *wts*^*X1*^ wing tumors and larvae with *scrib*^*1*^ *wts*^*X1*^ tumors additionally expressing Dome^ΔCYT^ in the PG under *phm-GAL4* control. Each dot plots mean GFP intensity in the PG measured in images like those in panel D. Horizontal lines represent mean values. Significance of differences in statistical tests is reported. Conducted tests were two-tailed t-test with Welch’s correction (control vs. *scrib*^*1*^ *wts*^*X1*^ tumor) and Mann-Whitney test (*scrib*^*1*^ *wts*^*X1*^ tumor vs. *scrib*^*1*^ *wts*^*X1*^ tumor expressing Dome^ΔCYT^ in the PG). (F) Pupation time in larvae with *scrib*^*1*^ *wts*^*X1*^ wing tumors and larvae with *scrib*^*1*^ *wts*^*X1*^ tumors additionally expressing Dome^ΔCYT^ in the PG under *phm-GAL4* control. Four repeats of the experiment are shown. Dot-connecting lines plot the accumulated percentage of pupated larvae through time. (G) Pupation time in *phm-GAL4* control animals and in both *phm-GAL4* and *phm>dome*^*ΔCYT*^ animals heated at 39^0^C for 2.5 hours during the mid L3 stage. Dot-connecting lines plot accumulated percentage of pupated larvae through time.

### JAK/STAT activation upregulates Apontic in the PG

To further investigate the action of JAK/STAT in the ring gland, we searched for known targets of JAK/STAT signaling that were expressed in the PG. We found one such candidate in Apontic, highly present in the PG according to published images (Rodrigues et al., 2021). Apontic (Apt), also known as Tracheae defective, is a transcription factor known to act as a downstream target of JAK/STAT in border cell migration (Starz-Gaiano et al., 2008). It is also involved in the development of multiple tissues, including the tracheal system (Eulenberg and Schuh, 1997). Staining with an anti-Apt antibody (Liu et al., 2014) confirmed expression of Apt in the PG and the entire ring gland (Fig. 6A). Furthermore, expression of Apt was highly upregulated in the PG in larvae containing *scrib wts* wing disc tumors and upon overexpression of Upd in the PG (Fig. 6A, quantified in Fig. 6B), while levels in tracheae associated with the ring gland were not affected. Apontic levels in the PG, in addition, were reduced upon *Stat92E* knock down in unchallenged larvae, and also in larvae with tumors upon dominant negative Dome^ΔCYT^ expression (Fig. 6A and B). All these results indicate that Apt expression is under JAK/STAT regulation in the PG.

**Figure 6.**
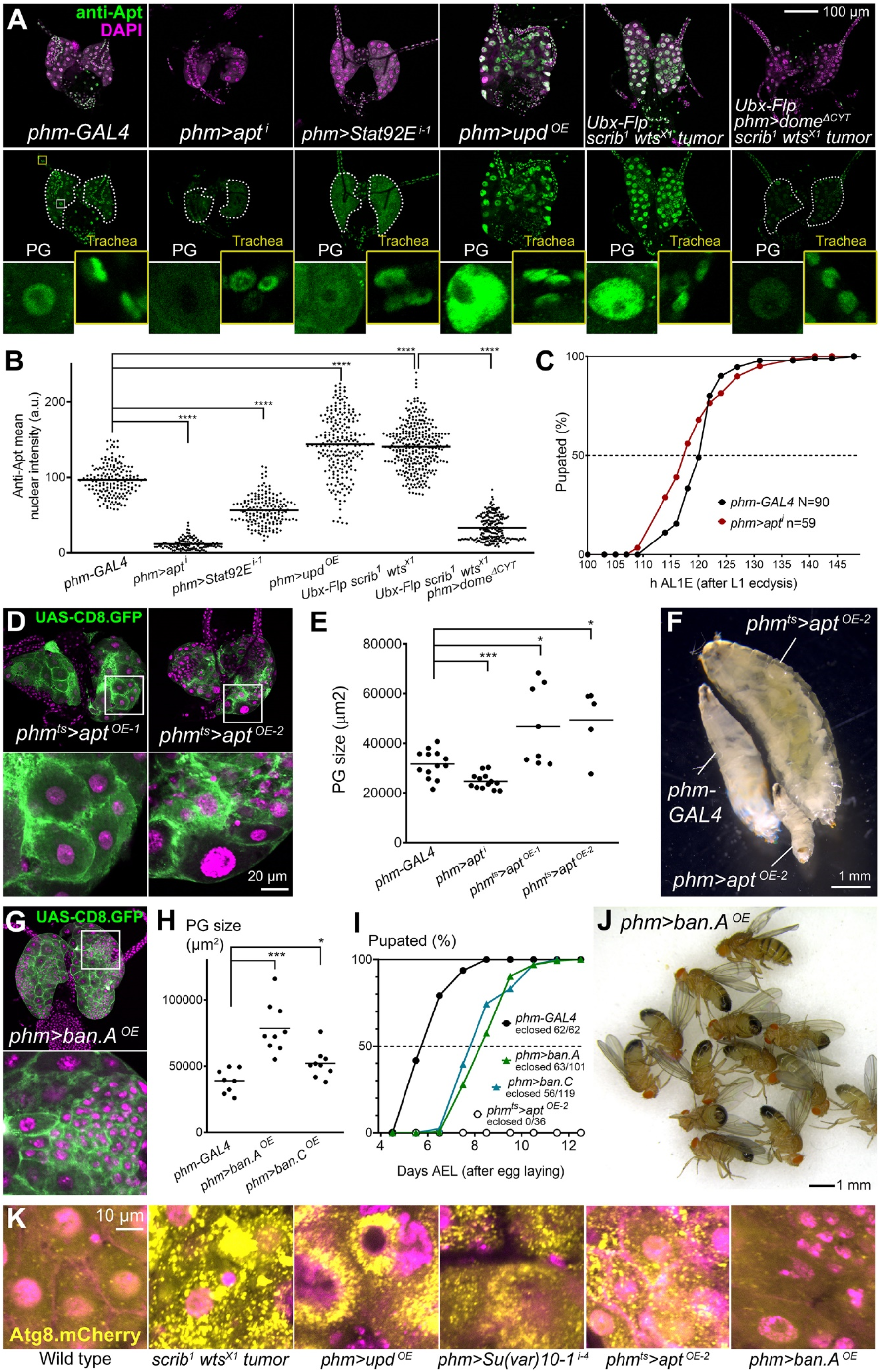
JAK/STAT target Apontic causes PG hypertrophy, autophagy and metamorphosis delay. (A) Anti-Apt antibody stainings (green, separate channel in lower panels) of wL3 ring glands of the indicated genotypes. Dotted lines represent PG outline. Nuclei stained with DAPI (magenta). Higher magnification insets show Apt expression in PG nuclei and in tracheal nuclei as an internal control where levels do not change. (B) Quantification of anti-Apt signal in wL3 PG from larvae of the indicated genotypes. Each dot plots mean GFP intensity in a PG nucleus measured in images like those in panel A. Horizontal lines represent mean values. n=178, 122, 173, 226, 320 and 202 nuclei, respectively. Significance of differences in statistical tests is reported. Conducted tests were Mann-Whitney tests except for *phm-GAL4* control vs. *phm>upd*^*OE*^ and *scrib*^*1*^ *wts*^*X1*^ tumors (two-tailed t-tests with Welch’s correction). (C) Pupation time in *phm-GAL4* control and *phm>apt*^*i*^ animals. Dot-connecting lines plot accumulated percentage of pupated larvae through time. (D) Ring glands from L3 giant larvae overexpressing *apt* in the PG under control of *phm-GAL4* and *tub-GAL80*^*ts*^. *phm-GAL4*-driven CD8.GFP in green. Animals were transferred at L2 stage from 18°C to 30^0^C to initiate *apt* overexpression. Overexpression of *apt* in *phm*^*ts*^*>apt*^*OE-1*^ and *phm*^*ts*^*>apt*^*OE-2*^ employs different transgenes (see Table S1). Nuclei stained with DAPI (magenta). Higher magnification insets (white squares) are shown in lower panels. (E) Quantification of PG size in control *phm-GAL4* wL3 larvae, *phm>apt*^*i*^ wL3 larvae and giant *phm*^*ts*^*>apt*^*OE*^ larvae. Each dot plots PG size measured in images like those in panels A and D. Horizontal lines represent mean values. Significance of differences in statistical tests is reported. Conducted tests were two-tailed t-test (*phm>apt*^*i*^), two-tailed t-test with Welch’s correction (*phm*^*ts*^*>apt*^*OE-1*^) and Mann-Whitney test (*phm*^*ts*^*>apt*^*OE-2*^). (F) Control *phm-GAL4* wL3 larva, *phm>apt*^*OE-2*^ L2 larva and giant *phm*^*ts*^*>apt*^*OE-2*^ L3 larva (L2 temperature shift). (G) Ring gland from a wL3 larva overexpressing miRNA *bantam* (*ban*) in the PG under *phm-GAL4* control. *phm-GAL4*-driven CD8.GFP in green. Nuclei stained with DAPI (magenta). Higher magnification inset (white squares) in lower panel. (H) Quantification of PG size in control *phm-GAL4* wL3 larvae and larvae overexpressing *ban* under *phm-GAL4* control. Each dot plots PG size measured from images like those in panel G. Horizontal lines represent mean values. Significance of two-tailed t-tests with Welch’s correction are reported. (I) Pupation time in *phm-GAL4* control, *phm>ban*^*OE*^ and *phm*^*ts*^*>apt*^*OE*^ animals. Dot-connecting lines plot the accumulated percentage of pupated larvae through time. (J) Despite metamorphosis delay, *phm>ban*^*OE*^ animals reach adulthood, unlike *phm*^*ts*^*>apt*^*OE*^animals. (K) Autophagy marker Atg8.mCherry (yellow) in the PG of wL3 larvae from the indicated genotypes. Nuclei stained with DAPI (magenta).

To test the effect of Apt upregulation, we overexpressed Apt in the PG. Apt overexpression under control of *phm-GAL4* produced larvae that did not progress beyond the L2 stage. However, when expression was restricted to the L2 and L3 stages by using the GAL80ts system, we observed hypertrophied PGs with enlarged nuclei (Fig. 6D and E) and inhibition of the larva/pupa transition (Fig. 6F), similar to the effect of JAK/STAT hyperactivation. Apt knock down, in contrast, produced slight reduction of average PG size and pupation time compared to controls (Fig. 6C and E). All these results suggest that Apt may be a target mediating the effect of JAK/STAT signaling both normal development and in response to tissue damage. A recent study, published while this manuscript was in preparation, found microRNA *bantam* (*ban*) was upregulated downstream of JAK/STAT signaling in the PG (Romão et al., 2021). We checked the effect of *ban* overexpression in the PG and found that it could also induce PG overgrowth (Fig. 6G and H). However, contrary to Apt expression, *ban*-overexpressing PGs contained regions with large numbers of apparently diploid cells, suggesting that *ban* upregulation can increase number of PG cells before polyploidization and/or induce depolyploidization. Also in contrast with Apt overexpression, flies overexpressing *ban* in the PG were capable of pupating and producing adults, while Apt-overexpressing flies did not pupate (Fig. 6I and J). Finally, we found that the PG in larvae with *scrib wts* wing disc tumors, similar to Upd overexpression, *Su(var)2-10* knock down and Apt overexpression, presents high levels of autophagy induction, as evidenced by formation of numerous large Atg.mCherry-positive vesicles in the cytoplasm, consistent with the vacuolation and varying degrees of tissue degeneration in these PG, and contrary to what is observed in the *ban*-overexpressing PG (Fig. 6K). Our results, therefore, suggest that different JAK/STAT targets may contribute to different aspects of the response to tissue damage (Fig. 7) and leaves for future studies a thorough analysis of these targets, their effects on PG development and their integration with other signals regulating developmental timing.

**Figure 7.**
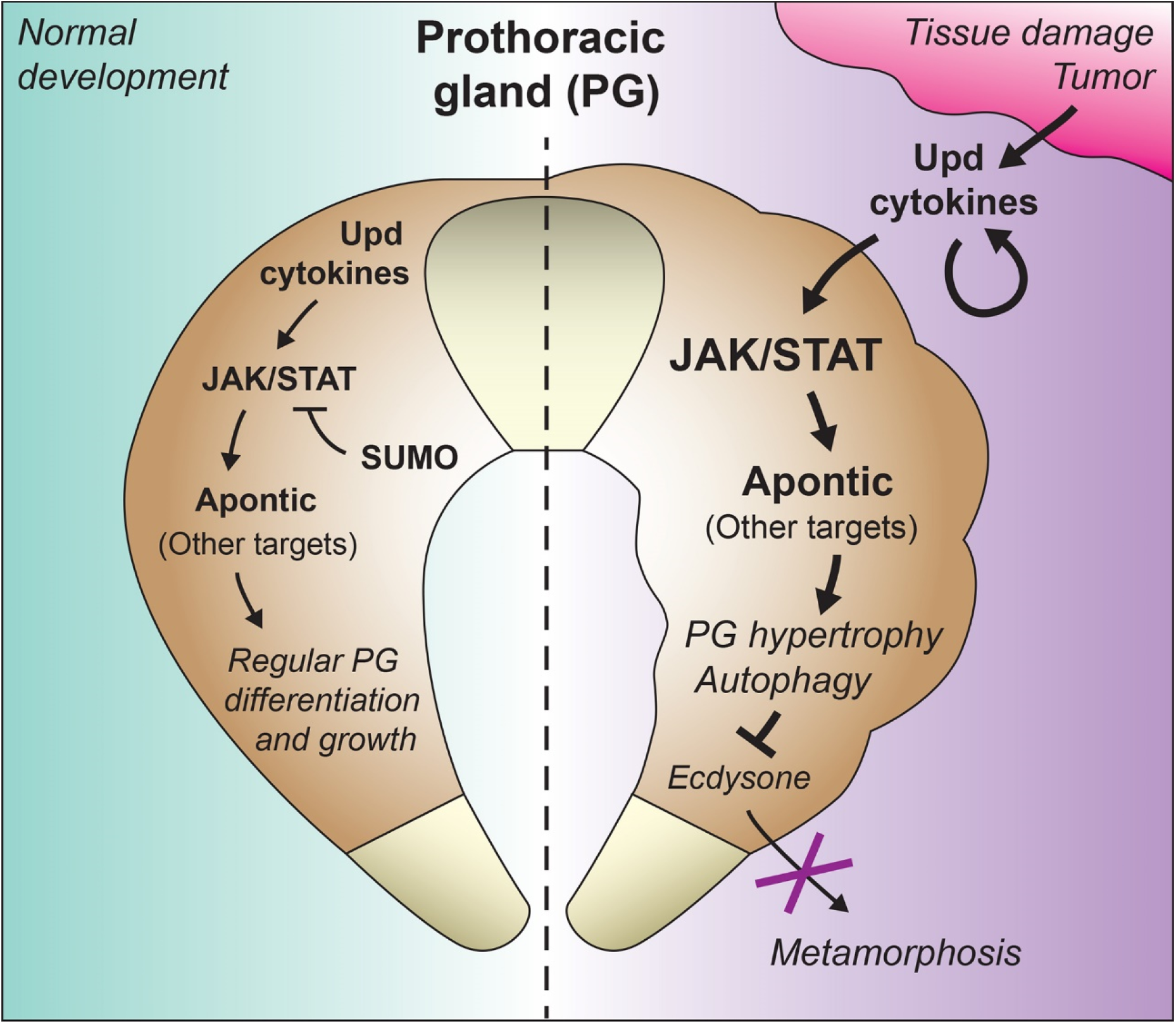
Intrinsic and extrinsic regulation of PG growth and developmental timing by JAK/STAT. Schematic model of the function of JAK/STAT signaling in the PG. Intrinsic expression of Upd3 in the PG activates basal levels of JAK/STAT activity in an autocrine way. Basal JAK/STAT activity is required for correct development of the PG, its loss reducing PG size and slightly advancing the onset of metamorphosis (larva/pupa transition). Levels of JAK/STAT activity are controlled through SUMOylation, involving SUMO E3 ligase Su(var)2-10 (PIAS), thus preventing excessive pathway activation. In conditions producing tumors and tissue damage, Upd cytokines secreted from the damaged tissues and amplified by other tissues (Pastor-Pareja et al., 2008) activate JAK/STAT signaling in the PG, which increases expression of JAK/STAT target Apontic and contributes to strong autophagy induction and inhibition of metamorphosis.

## DISCUSSION

Our results show that JAK/STAT signaling is active in the PG at low levels during normal development. Another study examined JAK/STAT activity in the PG and found in it as well expression of the same 10xSTAT-GFP reporter we used (Pan and O’Connor, 2020). We found, in addition, that pupation is slightly advanced and the size of the ring gland is smaller in multiple conditions of JAK/STAT signaling reduction, including knock down in PG cells of *upd3*, which a GAL4 expression reporter shows is intrinsically expressed in the ring gland. JAK/STAT signaling during embryonic development is known to be involved in the specification of the tracheal primordium that gives rise to the PG and corpus allatum (Sanchez-Higueras et al., 2014). Our results may therefore reveal a continued role of JAK/STAT for the development and differentiation of the larval PG.

Consistent with a role of JAK/STAT in ring gland development, hyperactivation of the pathway exhibited contrary effects, producing PG overgrowth and metamorphosis delay. PG overgrowth and ploidy increase is observed in conditions hyperactivating TOR/insulin signaling (Mirth et al., 2005; Ohhara et al., 2017). However, those conditions result in advanced pupation. This has been interpreted as revealing a mechanism by which insulin-driven PG growth mirrors organismal growth and triggers metamorphosis in well-fed animals. PG overgrowth resulting from JAK/STAT activation, in stark contrast, inhibits transcription of enzymes in the ecdysone synthesis pathway and results in absent or delayed metamorphosis. Also in contrast, JAK/STAT-induced PG hypertrophy leads to a highly aberrant tissue where vacuolation, autophagy and large differences in cell size are observed. We found in the literature that loss of *smt3*, encoding fly SUMO, caused a strikingly similar PG hypertrophy phenotype (Talamillo et al., 2008). Our results, showing that *Stat92E* knock down suppresses the effect of SUMO E3 ligase Su(var)2-10 knock down, indicate that hyperactivation of JAK/STAT signaling is responsible for this phenotype. This suppression result, in fact, further confirms a role of JAK/STAT in PG development, since PIAS has been shown to specifically bind and inhibit only active phosphorylated dimeric STAT, not monomeric STAT (Liao et al., 2000), and thus pathway hyperactivation in this condition strongly implies pre-existing activity.

In addition to its developmental role, JAK/STAT signaling is activated in the PG in response to tumors and tissue damage. Strong local upregulation of all three Upd cytokines has been observed in tumors (Wu et al., 2010), while amplification through systemic upregulation has been observed for Upd3 at least (Pastor-Pareja et al., 2008). Although not investigated here, our results predict that JAK-STAT activation by infection may regulate developmental timing as well, given the involvement of Upd cytokines in the immune response to pathogens (Agaisse and Perrimon, 2004). A study postulated that JAK/STAT signaling influences developmental timing by enhancing local expression of Dilp8 in wounded tissue (Katsuyama et al., 2015). Our results show that Upd cytokines, in addition, have a strong unmediated effect in the PG, where they activate JAK/STAT signaling and influence the larva/pupa transition directly. Partial rescue of tumor-induced pupation delay by expressing a dominant negative version of the JAK/STAT receptor Domeless in the PG clearly shows that JAK/STAT acts directly in the PG without mediation of Dilp8. Pupation delays induced by tissue damage are only partially rescued in null mutants for *Lgr3* (Garelli et al., 2015), encoding the specific receptor of Dilp8, which suggests that JAK/STAT, Dilp8 and perhaps other signals may be responsible for damage-induced pupation delay in parallel.

A number of pathway ligands acting in tissues in typical developmental patterning and differentiation/growth capacities have been shown to regulate ecdysone production by the PG. Such is the case of Hedgehog, released from enterocytes under starvation conditions (Rodenfels et al., 2014). TGFβ homolog Dpp, normally escaping from imaginal discs to the hemolymph (Ma et al., 2017), has been shown to downregulate ecdysone production by increasing nuclear FOXO localization and *ban* expression, suggesting that disc-derived Dpp functions as a signal that conveys organismal growth status to the endocrine system (Setiawan et al., 2018). Because Upd cytokines are produced during normal development in multiple organs, including imaginal discs, it is possible that PG-extrinsic Upd expression also serves a coordination role in normal development similar to Dpp. More recently, EGF ligands Spitz and Vein expressed intrinsically in the PG have been shown to regulate pupation as well, in this case stimulating ecdysone production (Cruz et al., 2020). This raises the converse question of whether EGF produced by other tissues can influence timing of the larva/pupa transition. Altogether, these studies paint a scenario in which the distinction between short-range developmental pathway ligands, long-range inter-organ communication hormones and immune response cytokines becomes increasingly blurred.

While we were finishing our study, the group of Marco Milan reported that Upd3 signals from the tumor to the PG to delay metamorphosis (Romão et al., 2021), consistent with our findings. According to that study, JAK/STAT inhibits pupation by inducing expression of the *ban* miRNA. Our study confirms the ability of Upd cytokines from damaged tissue to delay pupation while, in addition, showing that Apontic is a target of JAK/STAT signaling in the PG. The effects of Apontic, indeed, closely resembled those of JAK/STAT hyperactivation achieved through Upd overexpression or *Su(var)2-10* knock down, including tissue hypertrophy and autophagy. The response to JAK/STAT in the PG, therefore, is likely to involve multiple targets. Future studies should clarify the targets and mechanisms by which JAK/STAT signaling regulates developmental timing by the PG.

## MATERIALS AND METHODS

### *Drosophila* strains and genetics

Standard fly husbandry techniques and genetic methodologies were used to assess segregation of transgenes in the progeny of crosses, construct intermediate strains and obtain flies of the desired genotypes for each experiment (Roote and Prokop, 2013). The GAL4-UAS binary expression system (Brand and Perrimon, 1993) was used to drive expression of UAS transgenes under control of GAL4 drivers *phm-GAL4* (PG) and *Cg-GAL4* (fat body). Flies were reared at 25°C on standard fly medium in all experiments, except for *apt* overexpression experiments using the GAL80^ts^ thermosensitive GAL4 repressor (McGuire et al., 2003), in which crosses to obtain *phm*^*ts*^*>apt*^*OE*^ animals were left for three days at 18°C to avoid early lethality before transferring to 30°C. Genotypes of flies in all experiments are detailed in Table S1. Original fly strains used in this study are listed in Table S2.

### Imaging and image analysis

Ring glands and wing discs were predissected in PBS by turning larvae inside out with fine tip forceps, fixed in PBS containing 4% PFA (paraformaldehyde, Sinopharm Chemical Reagent, cat # 80096692), washed in PBS (3 × 10 min), dissected from the carcass and mounted on a glass slide with a drop of DAPI-Vectashield (Vector Laboratories, cat # H-1200). Confocal images were acquired with a Zeiss LSM780 confocal microscope equipped with Plan-Apochromat 20× NA 0.8 objective (ring gland images) and an EC Plan-Neofluar 10× NA 0.3 objective (wing disc images). Images of larvae, pupae and adults were taken with a Leica M125 stereomicroscope, except for the image in Fig. 4A, taken with a Leica MZ10F stereomicroscope.

For quantification of JAK/STAT activity in the PG, the polygon selection in ImageJ-FIJI software was used to outline the area occupied by the PG in confocal images of ring glands and mean 10xSTAT-GFP intensity (total fluorescence intensity divided by area) was measured inside. At least five ring glands were analysed in each condition.

For PG size quantification, the polygon selection in ImageJ-FIJI software was used to measure the area occupied by the PG in confocal images of ring glands. At least five ring glands were analyzed for each condition (each point represents the PG size in one ring gland).

For PG nuclear counts, nuclei were manually counted using the Multiple points tool in ImageJ-FIJI software. These counts were conducted on confocal Z-stack images of DAPI and CD8.GFP (expressed under PG-specific *phm-GAL4* control). At least 18 ring glands were analysed per genotype.

For ploidy estimation, ring glands from wL3 larvae were dissected, fixed and mounted in DAPI-Vectashield (Vector Labs). Z-stacks of images of DAPI and *phm-GAL4*-driven CD8.GFP were acquired in a Zeiss LSM780 confocal microscope equipped with a Plan-Apochromat 63X oil objective. On those stacks, the volume of PG nuclei was delimited and labelled with the Surface function in Imaris 9.3.1 software (Bitplane) and total DAPI fluorescence inside the nucleus recorded. Ploidy was calculated with reference to the average value in measurements of DAPI fluorescence conducted in the same way in diploid (2C) blood cells.

For quantification of anti-Apt signal, the polygon selection in ImageJ-FIJI software was used to outline the PG nuclei in confocal images of stained ring glands and mean intensity (total fluorescence intensity divided by area) was measured inside. More than 120 PG nuclei from at least 8 ring glands were analysed for each genotype.

### Immunohistochemistry

Anti-Apt stainings were performed following standard procedures for imaginal discs. Briefly, wL3 larvae were pre-dissected in PBS by turning them inside-out with fine tip forceps, fixed in 4% PFA for 15 min, washed in PBS (3 × 15 min), blocked in PBT-BSA (PBS containing 0.1% Triton X-100 detergent, 1% BSA, and 250 mM NaCl), incubated overnight with primary anti-Apt in PBT-BSA at 4^0^C, washed in PBT-BSA (3 × 20 min), incubated for 2 h with anti-rabbit IgG conjugated to Alexa-555 (1:200, Life Technologies) in PBT-BSA at room temperature, and washed in PBT-BSA (3 × 10 min) and PBS (3 × 10 min). Stained ring glands were finally dissected and mounted on a slide with a drop of DAPI-Vectashield (Vector Labs).

### Pupation timing

Fresh virgin female and male flies were kept together in bottles 3 days for mating and then allowed to lay eggs onto grape juice agar plates for 2 h at 25°C. The next day, newly hatched first instar larvae (L1) were picked at 1 h intervals and transferred to standard medium vials on which the top layer of food had been ground with forceps. 20 to 40 larvae were deposited in each vial. The number of pupated animals was counted at 2 to 6 h intervals. For the experiment in Fig. 6I, egg laying took place during 1 day in regular food vials instead and pupation time was computed in days after egg laying (±0.5 days).

### Quantitative RT-PCR

Total RNA was extracted from cephalic complexes containing the ring gland using TRIzol reagent (ThermoFisher Scientific), treated with DNase (Promega) and used as a template for cDNA synthesis using iScript™ cDNA Synthesis Kits (BIO-RAD). RT-PCR reactions were performed using SYBR Green Supermix (BIO-RAD) in a CFX96 Real-Time PCR system (BIO-RAD). Expression values were normalized to *rpL23* transcript levels. Fold change with respect to the wild type control was calculated with Bio-Rad CFX Manager 3.1 software. Three separate biological replicates were performed for each experiment, each with three technical replicates. Primers used were:

rpL23-F: GACAACACCGGAGCCAAGAACC, rpL23-R: GTTTGCGCTGCCGAATAACCAC;

phm-F: GGATTTCTTTCGGCGCGATGTG, phm-R: TGCCTCAGTATCGAAAAGCCGT;

dib-F: TGCCCTCAATCCCTATCTGGTC, dib-R: ACAGGGTCTTCACACCCATCTC;

sad-F: CCGCATTCAGCAGTCAGTGG, sad-R: ACCTGCCGTGTACAAGGAGAG;

nvd-F: GGAAGCGTTGCTGACGACTGTG, nvd-R: TAAAGCCGTCCACTTCCTGCGA.

### Wounding assay

Wounding of the larval epidermis was performed on mid L3 larvae by puncturing at once dorsal and ventral epidermis near the posterior end of the larva with fine tip forceps. Wounding operations were perfomed on sylgard plates on clean, dried larvae, avoiding damage to the gut or other internal organs. Operated larvae were kept on a dry slide for 5 min to allow coagulation at the wound and prevent bleeding. Operated larvae were finally transferred to standard medium vials on which the top layer of food had been ground with forceps.

### Heating assay

Newly hatched L1 larvae were picked at 1 h intervals and transferred to standard food vials after 4 h egg laying in agar juice plates, placing 20 to 40 animals per vial. 4 days after L1 ecdysis, mid L3 larvae were picked and transferred into 3.5-cm plastic dishes filled with ground standard food for heat treatment. The dishes were sealed with parafilm and placed for 2.5 hours in a water bath at 25°C (control) or 39°C. After heating, larvae were kept at room temperature for 15 min and then placed back into standard food vials in which the top layer of food had been ground with forceps. Pupation time after L1 ecdysis was recorded.

### Quantification and statistical analysis

Statistical analysis and graphical representations were performed with Graphpad Prism software. Horizontal lines in all graphs represent average values (means). For statistical comparisons, unpaired two-tailed Student’s t tests were conducted when data passed both D’Agostino-Pearson normality tests and F-tests for equal variance. Unpaired two-tailed t-tests with Welch’s correction were used when data passed D’Agostino-Pearson normality tests, but not F-tests for equal variance. Finally, non-parametric Mann-Whitney tests were used when data did not pass D’Agostino-Pearson normality tests. Significance of statistical tests is reported in graphs as follows: p<0.0001 (****), p<0.001 (***), p<0.01 (**), p<0.05 (*), and p>0.05 (n.s., not significant).

## Supporting information

Table S1_Detailed genotypes

Table S2_Original_strains

## ACKNOWLEDGMENTS

We thank Francisco Martin, Tian Xu, Jian-Quan Ni, James Castelli-Gair Hombria, Zhouhua Li, Herve Agaisse, the Tsinghua Fly Center, Bloomington Drosophila Stock Center, Vienna Drosophila Research Center and Kyoto National Institute of Genetics for fly strains, and Qing-Xin Liu for anti-Apontic antibody. We also thank Marta Rojas for assistance with RNA extraction and qRT-PCR. This research was supported by grants 31771600 and 91854207 from the Natural Science Foundation of China (NSFC).

## SUPPLEMENTAL MATERIALS

### Table S1. Detailed genotypes

Genotypes of animals in all experiments, listed by figure.

### Table S2. Original strains

Strains used in the study and their origin.

